# Targeted sequencing of expanded tandem repeats: Identifying interruptions and errors

**DOI:** 10.64898/2026.07.23.739298

**Authors:** Aeverie E. R. Heuchan, Ruban Rex Peter Durairaj, Alys N. Aston, Emma L. Randall, Laura Heraty, Christopher Smith, Alysha S. Taylor, Thomas H. Massey, Stephanie Tomé, Vincent Dion

## Abstract

Expanded repeat disorders remain without a disease modifying treatment. Some of the most important modifiers of disease severity include inherited repeat size, their somatic mosaicism, and whether they contain interruptions. Targeted sequencing approaches are becoming the gold standard on how these parameters are measured despite repeats being notoriously challenging to sequence. Here we developed CHARLIE (Comprehensive High-throughput Analysis of Repeat Length, Interruptions and Expansions) to identify and position interruptions within expanded repeats. We used samples from Huntington’s disease and myotonic dystrophy type 1 sequenced with Illumina MiSeq and PacBio’s Single-Molecule Real-Time Sequencing. We validated the pipeline against previous methods for identifying germline-inherited interruptions. One challenge that can potentially mask the identification of true interruptions and sequence variation is the accuracy of sequencing platforms, which, when applied to expanded repeats, is largely unknown. Our results suggest that each sequencing platform produces a distinct error profile and we show that PCR-free library preparation for SMRT sequencing improves sequencing accuracy. CHARLIE, therefore, provides a method that can help differentiate between trivial sequencing errors and those interruptions that modify disease presentation, which can be inherited or somatic in origin.

## Introduction

Over 60 human diseases are caused by the expansion of tandem repeats and include many neurological, neuromuscular, and neurodegenerative diseases (1, 2). One of the best characterised examples of this family of diseases is Huntington’s Disease (HD), where repeat expansion beyond 35 CAGs at the *HTT* locus leads to the disease, with full penetrance requiring 40 CAGs or more (3). The size of the inherited repeat determines over 60% of the variation in the age at disease onset (4, 5). Somatic expansion, the ongoing increase in repeat size within affected tissues, is thought to accelerate disease progression (6–9). The most common allele in HD contains an interrupting CAA motif in the penultimate position of the repeat tract. When duplicated, this interrupted allele correlates with a 10-year delay in disease onset, whereas the loss of the CAA motif correlates with an earlier onset and a more severe disease (6, 10–12). CAA interruptions are thought to slow somatic expansion, providing an attractive model for their contribution to disease presentation (11, 13, 14). Similarly, in myotonic dystrophy type 1 (DM1), which is caused by the expansion of a CTG repeat at the *DMPK* locus (15–17), CCG interruptions are frequent and modify disease presentation and severity (18, 19). Therefore, measuring repeat size, repeat size distribution, and interruptions is of crucial importance for diagnostics, patient management, and to support the development of therapeutics that target the repeat tract (13, 20).

Current diagnostic tests only provide modal repeat size, but novel technologies for sequencing expanded repeats have emerged that could address this shortfall (21–24). Whole genome sequencing using Illumina short read technologies can provide modal repeat size and interruptions at scale in cohorts of patients, but it is done at the expense of knowing the size distribution, or indeed, any somatic mosaicism in interruptions that may be present (25, 26). One way around this problem is to use targeted sequencing approaches. Illumina MiSeq with its 300 bp read size limit has been extensively used for HD with good results (27, 28). Using this workflow, somatic instability can be quantified and interruptions mapped - if they are known. However, some repeat loci, including in DM1, are too long for MiSeq and require long-read sequencing approaches, including Oxford Nanopore Technology’s (ONT) and PacBio’s Single-molecule Real Time (SMRT) long read sequencing. These technologies are revolutionizing the way in which expanded repeats are sequenced. ONT’s advantage is its low cost, but it is highly error-prone with a probability of an error at any one position being up to 45% within short tandem repeats of non-pathogenic sizes (29). In the case of ONT, the exact chemistry and base callers influence the apparent errors obtained (30). On non-repetitive sequences, high error rates are easily circumvented by having multiple reads covering the same locus. Sequencing errors are expected to be randomly distributed within each read and can be averaged out when determining a consensus sequence for a sample. This works well for identifying repeat modal sizes and interruptions that are inherited and thus in most cells [e.g., (31)]. However, it comes at the cost of averaging out any somatic mosaicism.

A more accurate long-read sequencing approach is SMRT sequencing. It works by sequencing the same molecule multiple times, rather than using different reads from the same sample, and producing a circular consensus sequence. By definition HiFi reads are those circular consensus sequences that contain errors at a frequency of 0.1% at any one base pair (21, 22). This higher accuracy means that it becomes possible to interrogate somatic mosaicism and begin to differentiate between interruptions of biological origins - known to impact phenotypes - and simple sequencing errors.

To differentiate between systematic sequencing errors and *bona fide* interruptions, it is critical to know how accurate the sequencing platforms are within expanded repeats. However, this information is lacking. Only one study, published in 2013 (32), addresses this question, albeit with an older SMRT chemistry, sequencing platform, and data processing pipeline. They showed that sequencing accuracy within the CGG repeat at the *FMR1* locus, responsible for Fragile X syndrome, was markedly lower than in the flanking sequences (32). Whether the same is true for CAG/CTG repeat loci and with newer technologies is not known.

Here we have developed CHARLIE (Comprehensive High-throughput Analysis of Repeat Length, Interruptions and Expansions). CHARLIE determines the presence and frequency of interruptions within expanded repeats and positions the interruptions within the repeat tract unsupervised. CHARLIE is agnostic to the sequencing platform used or the exact expanded repeat motif, making it broadly applicable. Importantly, it allows the determination of interruptions at the single read level.

## Results

### CHARLIE

CHARLIE is designed to find interrupting motifs peppered within repeats and to find their position within the repeat. The ‘inter find’ mode (Figure 1A) of CHARLIE uses a FASTX file as input and the user-defined sequence of the repeated motif (e.g. CAG). It then searches iteratively for every possible combination of interrupting bases. To ensure searches are targeting interruptions of the repeat tract, and not identical sequences in flanking regions, one (or more) repeat motif(s) is placed either side of the interruption in each search query (see specificity in the Methods section). For instance, a specificity of 1 looks for interruptions within a CAG repeat using (CAG)_1_- interruption-(CAG)_1_ as a search query, whereas a specificity of 2 searches (CAG)_2_- interruption-(CAG)_2_. CHARLIE returns the frequency of each interruption occurring at least once in a read. The size of the interrupting motif is user-defined and here we used interruptions of mono- di- and trinucleotides.

**Figure 1.**
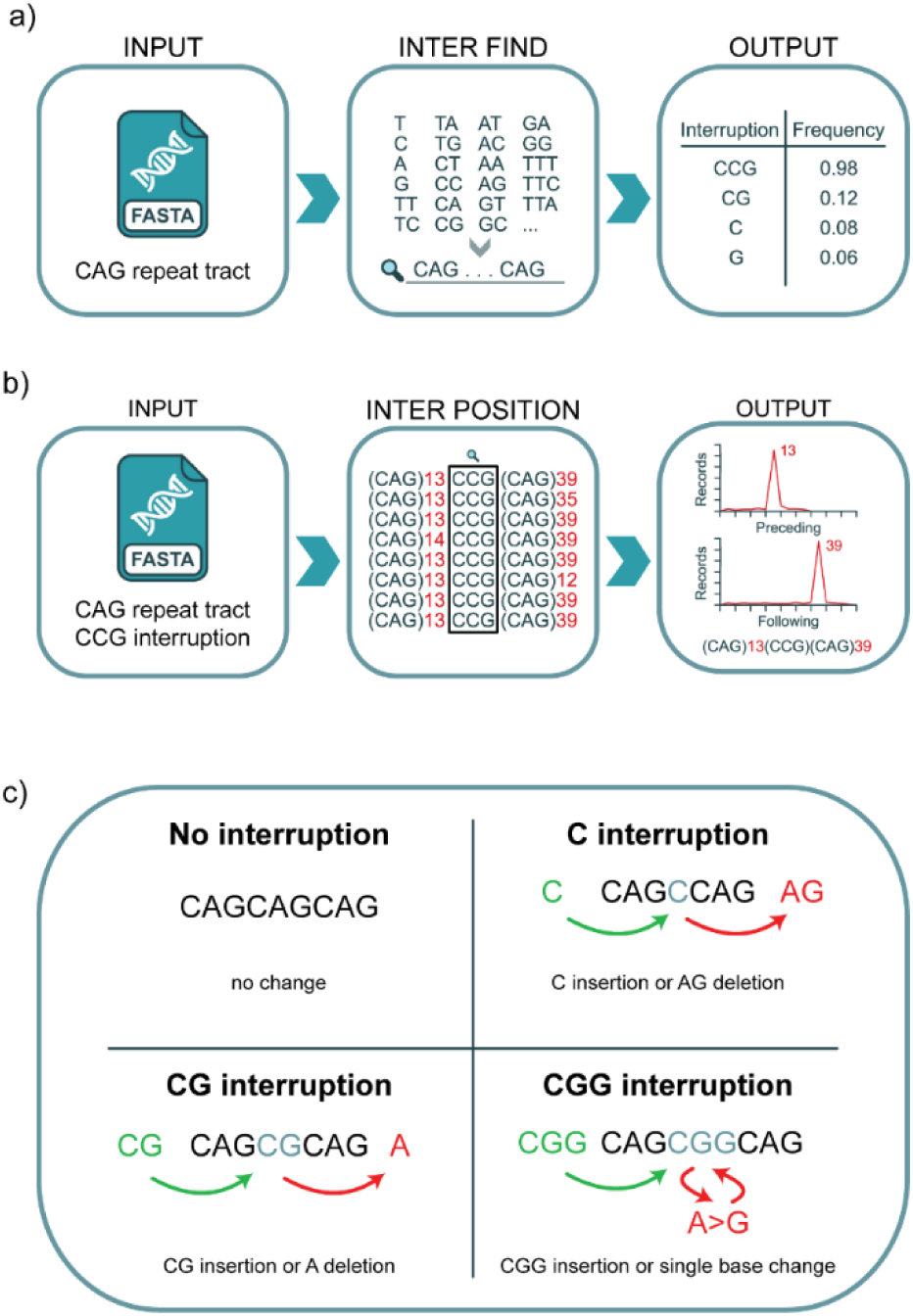
CHARLIE. a) Inter find takes a FASTX file of the DNA sequence as input and searches for every possible interruption up to n bases in size. Outputs are reported as frequency of reads containing each interruption. b) Inter position takes a FASTX file of the DNA sequence, alongside a previously identified interruption as input. The number of repeat motifs between the interruption and the beginning or end of the repeat tract is separately calculated. Outputs are reported as the number of reads at each number of preceding/following repeat motifs. From these outputs a modal interrupted repeat sequence is determined. c) Examples of reported 1-3 base interruptions and the mechanisms that could be responsible for them.

The ‘inter position’ mode (Figure 1B) uses the same FASTX as input to determine the most common number of repeats flanking an interruption within the reads of a sample, thereby positioning the interruption.

CHARLIE is blind to the mechanism that creates any interruption it reports (Figure 1C). For example, a C interruption in a CAG repeat tract could be caused by the insertion of a single C nucleotide, or the deletion of an AG dinucleotide. Moreover, it is blind to whether the sequence change is of biological origin (germline or somatic) or from a simple sequencing error. So as to not suggest origin or mechanism, hereafter we will use the term interruption to refer to any deviation from the pure repeat sequence.

We benchmarked CHARLIE against a well-defined dataset (10) composed of 641 blood and lymphoblastoid samples from HD patients, the HD MiSeq dataset, which was analysed previously with both ScaleHD (33) and Repeat Detector (27). In this dataset, the previously determined number of samples with an expanded allele containing at least one CAACAG motif was 580 (90.5%, Supplementary Table 1). We used CHARLIE to search the same dataset and found the same 580 samples containing alleles with the CAGCAACAG motif. In these samples, at least 90% of the reads contained the interruption (Supplementary Table 1). By contrast, among the remaining 61 samples, the frequency of reads with this interruption remained below 10%, suggesting that there were no samples with an ambiguous call (Supplementary Table 1).

One sample in this dataset is known to contain a CAC interruption on the expanded allele. This interrupted allele was first found manually and then added as a known interruption in the ScaleHD library of reference sequences (10). Repeat Detector (27) identified the presence of the interruption in this sample, but the motif again had to be identified manually. CHARLIE was designed to identify the presence of an interruption, its motif, and position of the interruption within the repeat tract unsupervised. And indeed, CHARLIE readily identified the interruption in this sample with 94.9% of the reads from the expanded allele having at least one CAC interruption (search motif: CAGCAGCACCAGCAG, Supplementary Table 1).

If this interruption is of germline origin, we expect that most reads will contain the interruption, as is the case here, and whilst the number of CAG repeats 5’ and 3’ of the interruption will be similar between reads, we expect some variation due to somatic instability. By contrast, a sequencing error is expected to occur randomly across the repeat tract and thus be poorly positioned by CHARLIE. In this case, CHARLIE found that the most frequent structure of the CAC-containing allele is (CAG)_41_CAC(CAG)_3_ with 32.9% of interrupted records having this exact structure (Supplementary Figure 1, Supplementary Table 1). This is the same structure that was determined previously (10). To determine whether the interruption was biological and not due to a generally high level of sequencing errors in this sample, we looked at all other possible interrupting motifs. The most common was CGG, found in 16% of the reads. Despite being a commonly identified interruption, ‘inter position’ found little evidence of a positioned CGG interruption with the most frequent allele structure being (CAG)_1_CGG(CAG)_1_ found in 2.4% of the interrupted alleles, or in 0.02% of all reads (Supplementary Figure 1), suggesting sequencing errors rather than an interruption of a biological origin. By contrast, the CAC interruption is normally rare. Indeed, of the remaining 640 samples, the highest frequency of reads containing a CAC within a sample was 1.3% and it was poorly positioned (Supplementary Table 1). Taken together, these results suggest that CHARLIE accurately identifies and locates interruptions that are inherited and thus present in most reads. Unlike other methods, it does so without prior knowledge of the interruption or its position.

### Error frequencies are similar within the repeat tract and in the flanking sequences

Interruptions seen at higher frequencies, of a different type than commonly seen in a given sequencing platform, and at a specific position within the repeat tract are more likely to be of biological origin. CHARLIE is ideally suited to identify these characteristics and here we applied it to assessing the types and frequency of sequencing errors made by SMRT sequencing and MiSeq within the expanded repeats.

In both cases we only used the high-quality reads, which are those that are expected to contain a probability of an error at anyone base pair of 0.1% or better (MiSeq Q30 and SMRT HiFi). We left MinION datasets out of this analysis because the error rates are too high (up to 45% for any given position (29)) and therefore nearly all reads are expected to have at least one interruption given the length of the repeats (27, 29).

We used CHARLIE on several datasets: the HD MiSeq dataset described above; the PCR-based HD SMRT dataset consisting of 17 HD samples characterised in depth previously (27); a new PCR-free HD PureTarget dataset of 4 samples from a HD patient-induced pluripotent stem cell (iPSC) line; and two DM1 SMRT samples each consisting of two independently sequenced samples from a mother/daughter pair with the same interruption and a similar repeat size. Both samples were previously sequenced, A3 without PCR and A4.1 with PCR (18, 34).

Assuming a Poisson distribution for interruptions due to sequencing errors and 0.1% error rates, we expected that a repeat tract of 40 triplet repeats (120 bp) would contain on average at least one interruption in 12% of the reads. This frequency of interruptions is expected to rise with the length of the read, which is determined here by the size of the repeat. The first observation that we made was that the frequency of interruptions was higher than what is expected from the error rates advertised by the companies. Indeed, in the HD MiSeq dataset, the average proportion of reads with at least one interruption was 30.9% ± 4.7% (Supplementary Table 2). This frequency increased with repeat size, as expected (Figure 2a). In the HD SMRT dataset, the frequency of reads with at least one interruption was similar, with 39.4% ± 7.3%, and increased with read size (Figure 2b, Supplementary Table 2). The difference was expected because the repeat sizes are longer in the HD SMRT dataset compared to the HD MiSeq dataset. Indeed, interruption frequencies were similar when comparing the repeat sizes (Figure 2ab).

**Figure 2:**
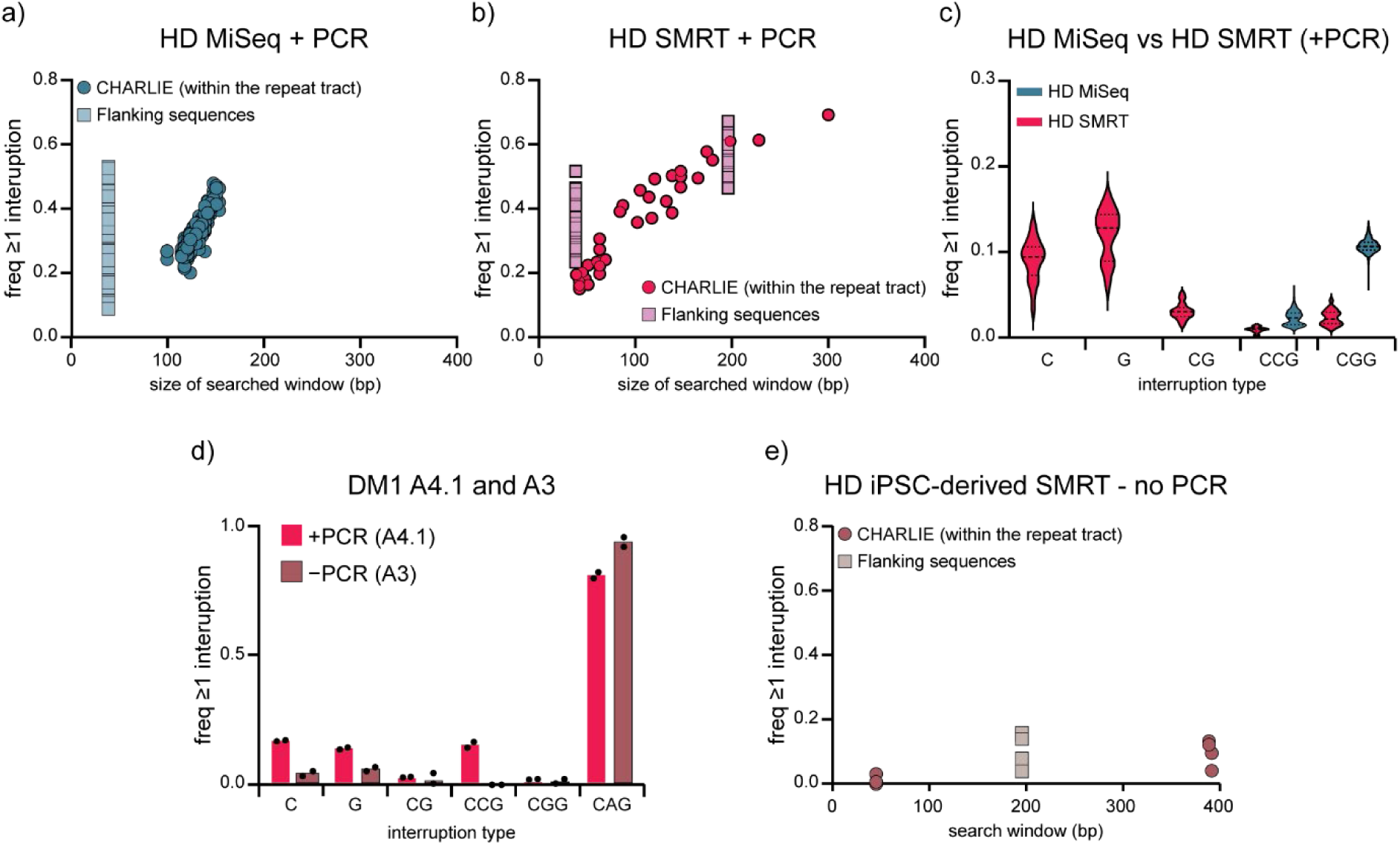
Interruption profiles in SMRT and MiSeq datasets. A) Frequency of reads with at least 1 interruption in the HD MiSeq dataset using CHARLIE (circles) or the 38 bp immediately upstream of the repeat tract (flanking – squares). n=641 – only the expanded alleles are shown. The full dataset is in Supplementary Table 1). The sample with the CAC interruption was removed from this graph for clarity as it returned an interruption frequency of 96.3% (see Supplementary Table 1). B) Frequency of reads with at least 1 interruption in the HD SMRT dataset (+PCR, n=17) using CHARLIE (circles) within the repeat or a 38bp or 196 bp window immediately upstream of the repeat tract using the flanking sequence pipeline (squares). C) Comparison of the type of interruptions made at the HD locus by SMRT (n=17) and HD MiSeq (n=641) datasets when using PCR in the library preparation. D) Comparison of the types of interruptions seen in the DM1 A4.1 and A3 datasets comparing library preparation methods that utilised PCR Cas9-based enrichment approach (A3; n=2) or a regular amplicon sequencing (A4.1; n=2). This sample contains a verified CAG interruption that is germline in origin. E) Frequency of reads with at least one interruption when HD iPSC (n=2) and HD iPSC-derived neurons (n=2) were sequenced using the PCR-free PureTarget kit from PacBio.

To understand whether these higher interruption frequencies are specific to the repeat tract, as observed with CGG repeats in 2013 (32), we analysed the interruption frequency in the non-repetitive sequence immediately upstream of the repeat tract (Supplementary Figure 2). This was done using a separate code since CHARLIE can only be applied to repetitive sequences. The flanking analysis used a reference sequence specific for the *HTT* locus (see methods), immediately upstream of the CAG repeat. This was done because downstream of the repeat tract there is a highly polymorphic CCG repeat (6, 10, 28), which would complicate the analysis.

In the HD MiSeq dataset, there are only 38 bp sequenced immediately upstream of the repeat tract to accommodate the limit on the read size for MiSeq, whereas in the HD SMRT there are 196 bp. We therefore compared the 38 bp immediately 5’ of the repeat tract in both datasets and used both 38bp and 196 bp in the HD SMRT dataset. We found that the interruption frequencies appear slightly higher for the non-repetitive upstream sequence compared to the repeat tract itself (Figure 2ab, Supplementary Table 3).

We then compared the interruption frequency between a search window of 38 bp and one closer to the size of an expanded allele, 196 bp, using the HD SMRT dataset. We found that, as expected, the frequency of interruptions increased with the size of the searched window. Moreover, there was no difference in the interruption frequency between the repeat tract and the upstream sequences when we considered this longer search window of 196 bp (Figure 2b, Supplementary Table 2). Taken together, we find no evidence for sequencing errors being higher within the repeat tract compared to the flanking sequences, contrary to what was found with older chemistries (32).

### Sequencing errors appear distinct between sequencing platforms

Because CHARLIE can identify the types of interruptions in addition to their frequencies, we next asked whether there were platform-specific interruptions, which is suggestive of sequencing errors rather than a change that is biological in nature. In the HD MiSeq dataset, the most common interruptions were CGG triplets with an average of 8.2% of the reads containing at least one within the repeat tract (Figure 2c, Supplementary Table 1). By contrast, the HD SMRT dataset contained high levels of C and G interruptions at 9.1% and 12%, respectively (Figure 2c, Supplementary Table 4). These different interruption profiles may originate from errors made during the PCR amplification prior to library preparation. Indeed, the two HD datasets were made with different DNA polymerases in different labs and at different times.

To tease this apart, we ran CHARLIE on the PCR-based DM1 sample, A4.1, which was sequenced independently twice using SMRT sequencing (18). This sample contains a known CAG interruption within its CTG repeat. CHARLIE identified it readily in 82% of the reads on average between the two independent sequencing runs from this sample. The positioning was consistent with both runs returning (CTG)_29_CAG(CTG)_105_ (Supplementary Table 5), consistent with prior work (18). When comparing the types of interruptions seen in this sample, we found them to be similar to those in the HD SMRT dataset, with C (17% of the reads) and G (14 % of the reads) interruptions being the most frequent (when excluding the CAG interruption of biological origin in this sample, Supplementary Table 5). CGG interruptions, often seen in the HD MiSeq datasets but not in the HD SMRT dataset, were also rare in the DM1 SMRT dataset (1.3% - Figure 2d, Supplementary Table 5). Despite the use of different DNA polymerases for library preparation, similar error profiles are found in the DM1 and HD SMRT datasets. These data suggest that some of the errors appear to be platform-specific.

### PCR-free library preparation improves sequencing accuracy

To determine what proportion of the interruptions seen in the sequencing data was due to PCR, we analysed the sequencing data from the A3 DM1 sample using a Cas9 enriched, amplification-free library preparation method (34). We used CHARLIE to analyse the repeat tract and found that the frequency of interruptions was reduced across different motifs (Figure 2d, Supplementary Table 5). The exception was the CAG interruption - of biological origin -, which increased in frequency from 81.9% of the HiFi reads having the interruption (in A4.1 with PCR) to 94.9% (in A3 without PCR). The algorithm found that the most frequent allele structure was (CTG)_29_CAG(CTG)_107_ for one of the runs and (CTG)_29_CAG(CTG)_106_ for the second (Supplementary Table 5). These results suggest that PCR-free methods may increase the accuracy of detecting inherited interruptions.

To confirm that a Cas9 enriched, amplification-free library preparation method is less error-prone, we used the newer PureTarget kit (version 1) for SMRT sequencing. This kit contains sgRNAs to isolate 20 repeat loci and sequence them without the need for PCR. We prepared PureTarget libraries from 4 samples from the same HD patient-derived iPSC line with 135 repeats and 19 repeats on the expanded and non-expanded allele, respectively (35). Two of these samples were independently differentiated into neurons before DNA extraction and sequencing. We filtered the HiFi reads using the PacBio TRGT pipeline (36) to identify haplotypes and ran CHARLIE independently on the two *HTT* alleles. We found the frequency of interruptions to be markedly lower at 5.4% (0.97% for the short allele and 9.9% for the long allele Supplementary Table 1). On the expanded allele, the two most common interruptions were CA (3.5% of the reads) and G (2.7% of the reads – Supplementary Table 4). The interruption frequency in the upstream sequence was also lower compared to PCR-based libraries (Figure 2e). Taken together, these results suggest that PCR errors contribute to the interruptions found.

## Discussion

Here we developed a novel algorithm, CHARLIE, that can identify interruptions and position them along a repeat tract unsupervised, something that has not been possible with other methods. It is designed to help us distinguish between trivial sequencing errors and interruptions of biological origin that may modify clinical presentation. We applied CHARLIE to determining the error profile within expanded CAG/CTG repeats of two commonly used sequencing platforms. Our results suggest that PCR-free library preparation followed by SMRT sequencing can improve the accuracy of the data generated and help identify interrupted alleles in expanded repeats that are of biological origin instead of simple sequencing errors.

A limitation of CHARLIE is that interruptions that are very close together can be missed. Moreover, if the same interruption is found multiple times on the same read, CHARLIE will count it only once. We have mitigated this limitation here by reporting the number of reads with at least one interruption rather than being able to report all the interruptions present.

The results presented will be of importance in distinguishing interrupted alleles from sequencing errors within patient samples and thereby help improve the predictive power of diagnostic tests and support the development of safe DNA-targeting therapies.

## MATERIAL AND METHODS

### Sequencing datasets

The HD MiSeq Dataset contains 649 samples (544 immortalised lymphoblastoid, 98 blood, 7 known controls) obtained as described (10, 27). Of these, we removed 8 samples that had lower quality sequences as before, giving a total of 641 samples. The HD SMRT Dataset contains sequences from 22 HD lymphoblastoid samples obtained from the Coriell BioRepository and described in (27), here also, we removed 5 samples that had lower quality sequences for this analysis. The samples are available at Gene Expression Omnibus (GSE199005). The DM1 dataset contains sequences from two DM1 blood samples obtained from related individuals (mother and daughter), referred to A.3 and A4.1 with the same interruption and similar repeat size. The samples were described previously (18, 34, 37).

### Cell culture

Induced pluripotent stem cells were seeded in plates coated with Corning Matrigel (according to manufacturer’s guidelines) and cultured in Gibco E8 Flex Medium at 37°C, 5% CO_2_. E8 medium was supplemented with 10 µM ROCK Inhibitor (RI) during passaging and removed within 24 hours. E8 media was replaced with fresh every 2-3 days. To split or pellet cells for DNA extraction, cells were dissociated from plates by gently washing with PBS, incubating with ReleSR (STEMCELL Technologies) and centrifuging cells at 300xg for 3 mins. Pellets were stored at –80°C until DNA extraction.

IPSCs were differentiated into cortical neurons using the protocol outlined in (38): 60-70% confluent iPSC were pre-treated for 1hr with E8 media supplemented with 10μM RI and then dissociated to single cells using Accutase, pelleted, and resuspended in E8 supplemented with RI. Cells were seeded to plates pre-treated with Matrigel and left overnight to adhere. The next day cells were washed with PBS, then maintained in SLI media (Gibco Advanced DMEM-F12 with 1% GlutaMAX, 1% PenStrep, 1% MACs Neurobrew w/o VitA, 10μm SB431542, 200nM LDN-193189 and 1.5μM IWR-1-endo) for 8-12 days. Media was replaced every day. On day 8-12, cells were treated with RI, split using Accutase, and plated in NB media supplemented with RI (Advanced DMEM-F12 with 1% GlutaMAX, 1% PenStrep and 2% MACs Neurobrew (w/o) Vitamin A). RI was removed from the media the following day and the media was changed daily until day 16 by which time the cells become neural precursor cells (NPCs). For terminal differentiation NPCs were plated on poly-L-lysine- and Matrigel-treated plates in SJA media (Advanced DMEM-F12 with 1% Glutamax, 1% PenStrep, 2% MACs Neurobrew with Vit A, 2μM PD0332991, 10μM DAPT, 10ng/mL BDNF, 10μM Forskolin, 3μM CHIR99021, 200μM Ascorbic Acid, 0.8 mM CaCl2, 300 μM GABA) supplemented with RI for 24hours. NPCs were maintained in this media for 7 days with half media changes every 2 days. Following this, cells were maintained in SJB medium (Advanced DMEM-F12 with 1% GlutaMAX, 1% PenStrep 2% MACs Neurobrew with Vit A, 2μM PD0332991, 200μM Ascorbic Acid, 0.4mM CaCl2) for a minimum of 14 days to allow for neuronal maturation.

### PacBio library preparation and sequencing

The HD SMRT and DM1 SMRT datasets were sequenced previously (18, 27, 34). For the PCR-free HD SMRT dataset from iPSCs and iPSC-derived neurons, genomic DNA was extracted with either the PanDNA NanoBind kit (Pacific Biosciences), or the Monarch HMW DNA extraction kit (New England Biolabs). The PureTarget (103-390-400) library preparation was done according to the manufacturer’s instructions (Pacific Biosciences). Library concentration was quantified using the Invitrogen Qubit HS dsDNA kit. Sample pools were sequenced using the Pacific Biosciences Sequel IIe. Supplementary Table 6 contains the number of reads in each sample.

### Repeat Sizing & Data Curation

#### Repeat Sizing

All repeat sizing hereafter referred to was carried out using Repeat Detector v1.0 (27) (available at: https://github.com/DionLab/RepeatDetector) with parameters ‘--with-revcomp’, ‘-o histogram’, and ‘--cycle-range [0:1000]’. All repeats were sized using the permissive profile using parameter ‘--prf RepeatDetector/Profiles/CAG/Annex2_cag.prf’. When determining repeat sizes for the purpose of filtering FASTA files by repeat size, log files were generated in Repeat Detector using parameters ‘--prf RepeatDetector/Profiles/CAG/Annex2_cag.prf’ and ‘--with-revcomp’. Outputs were sent to log files using ‘> <sample>.logs’ at the end of the command.

#### Filtering

All samples of the HD MiSeq dataset were filtered for a quality score of Q30 (99.9% accuracy) using the tool fastq-filter (available at: https://github.com/LUMC/fastq-filter) with parameter ‘--mean-quality 30’. Average pass rate was 46.0%. To remove incomplete reads and reads from wild type alleles from the HD MiSeq dataset, all samples were subsequently filtered to only contain reads with 36 or more repeats, selected due to pathogenicity beginning at 36 repeats (39) using Repeat Detector log files and filter_logs_miseq_36+.py. The output filtered log files were used to filter the FASTA files with the script filter_fasta_miseq_36+.py. Repeat sizing was re-run on the filtered samples and the redetermined modal repeat sizes used for graphing. All custom python scripts can be found on the DionLab Github (https://github.com/DionLab/CHARLIE and https://github.com/DionLab/flanking)

Samples of the HD SMRT dataset were separated into short and long reads using repeat size on a sample by sample basis based on Repeat Detector log files for permissive repeat size determination (see filter_logs_pacbio.py and filter_fasta_pacbio.py). After exclusions and separation of long and short alleles there were 37 usable samples. Repeat sizing was re-run on the filtered samples and the redetermined modal repeat sizes used for graphing. For the DM1 datasets, repeat size was determined using Repeat Detector with the permissive settings, and split for long reads (≥11 CTGs) using filter_logs_DM1.py and filter_fasta_DM1.py before being analysed with CHARLIE.

For the HD PureTarget dataset, the alleles in the samples were processed like the HD SMRT dataset to obtain the HiFi reads before being processed using CHARLIE. We omitted the sequence CAGCAGCTCAGCAG in the analysis because it was present only outside the repeat tract and outside the HD MiSeq and HD SMRT amplicons. The TRGT pipeline for PureTarget analysis usually includes both HiFi reads and failed reads for the analysis. Here we have only used the HiFi reads to be able to compare them to the HiFi reads from amplicon-based library preparation methods.

### Characterisation of Interruptions using CHARLIE

#### Inter find

All samples were run through CHARLIE inter find with parameters ‘--revcomp’ and ‘--threshold 0’. Samples were run once with parameter ‘--specificity 1’(spec1) and again with ‘--specificity 2’ (spec2). Interruption frequencies generated with specificity 2 were used in graphing as they exclude sequences outside the repeat tract more readily. For samples in the HD datasets the default maximum interruption size of 3 was used. For samples in the DM1 datasets, the parameter ‘--maximum 4’ was used. More than one interruption can occur on the same read and therefore the adding up the total frequencies for every interruption adds up to more than 100%. One sample, with the CAC interruption, was left out of Figure 2 because it contained a CAC interruption of biological origin observed before (10, 27) with a disproportionally high frequency compared to the rest of the samples.

#### Inter position

CHARLIE inter position was run for the CAC interruption within the sample with parameters ‘--revcomp’, and ‘--graphcrop’. As a negative control the CGG interruption from the same sample was run with the same parameters.

### Interruption frequencies within flanking sequences

To scripts to assess error frequencies in the flanking sequences are found at https://github.com/DionLab/Flanking. Fastq files were converted into fasta using seqkit tool (v2.10.0). The compare_sequence.py (Supplementary Figure 2) was used to define the length of the flanking sequence to be interrogated (----[upstream]CAG[>=6], ----CTG[>=6] [downstream]----). The script then extracts the defined number of bases (e.g., 196bp) from flanking regions of the 5’ and 3’ of the repeats. The script uses difflib to compare extracted segments to the user defined target sequence and the reverse complement. Another custom python script (csv_process.py) was then used to process all the resulting csv files and output the percentage of reads without errors and the total number of reads. 5.2% of the alleles appeared to have a single A to G nucleotide polymorphism 27 bp upstream of the repat tract. For the alleles that contained this allele, we adjusted the reference sequence before calculating the error rates.

## ACKNOWLEDGEMENTS

We thank Joanne Morgan for assistance with running and maintaining the Sequel IIe and members of the Dion lab for comments on the manuscript. The Registry study is supported by the European Huntington’s Disease Network (EHDN), funded by CHDI Foundation, Inc. The funding source had no role in study design; in the collection, analysis, and interpretation of data; in the writing of the report; and in the decision to submit the paper for publication. We acknowledge the support of the Supercomputing Wales project, which is part-funded by the European Regional Development Fund (ERDF) via the Welsh Government.

## AUTHOR CONTRIBUTIONS

AERH Conceptualization, Data curation, Formal analysis, Investigation, Methodology, Software, Visualization, Writing – original draft.

RRD: Formal analysis, Methodology, Writing – review and editing.

ANA: Investigation, Methodology, Writing – review and editing.

ELR: Investigation, Methodology, Writing – review and editing.

LH: Investigation, methodology, Writing – review and editing.

CS: Validation, Writing - review and editing.

AST: Supervision, Validation, Writing – review and editing.

TM: Resources, Writing - review and editing.

ST: Methodology, Resources, Writing – review and editing.

VD: Conceptualization, Formal analysis, Visualization, Funding acquisition, Methodology, Project administration, Supervision, Writing – review and editing.

## DATA AVAILABILITY

CHARLIE is available at https://github.com/DionLab/CHARLIE and the version used here is found here. The codes for the flanking sequences are held here: https://github.com/DionLab/flanking. The HD SMRT dataset is available here at the Gene Expression Omnibus (GSE19900) while the HD PureTarget dataset is available at https://www.ncbi.nlm.nih.gov/sra/PRJNA1484039.

---

**Supplementary Figure 1:**
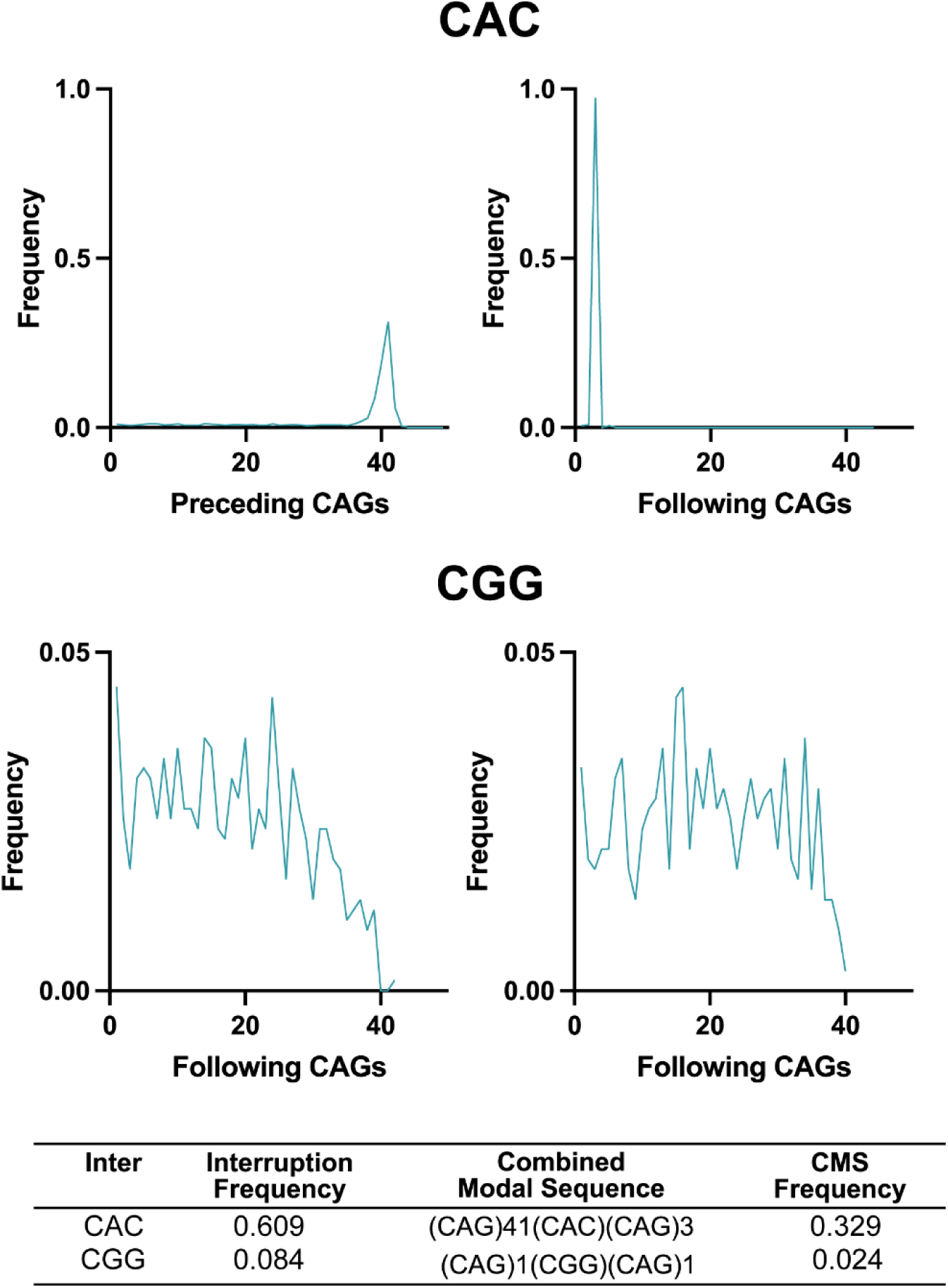
Inter position functionality of CHARLIE. A) Using CHARLIE’s Inter Find script, we identified the sole sample with a CAC interruption in the HD MiSeq dataset. This was found in the whole sample at a frequency of 0.609 and, when looking only at the expanded allele, the frequency of reads with the interruption was 0.949. The Inter position function allowed us to position it precisely within the repeat tract (Top). The most common structure was (CAG)_41_(CAC)(CAG)_3_ at a frequency of 0.329 of the interrupted reads. By contrast, CGG appears to be a sequencing error due to its low interruption frequency (0.084) and combined modal sequence (CMS) frequency (0.024).

**Supplementary Figure 2:**
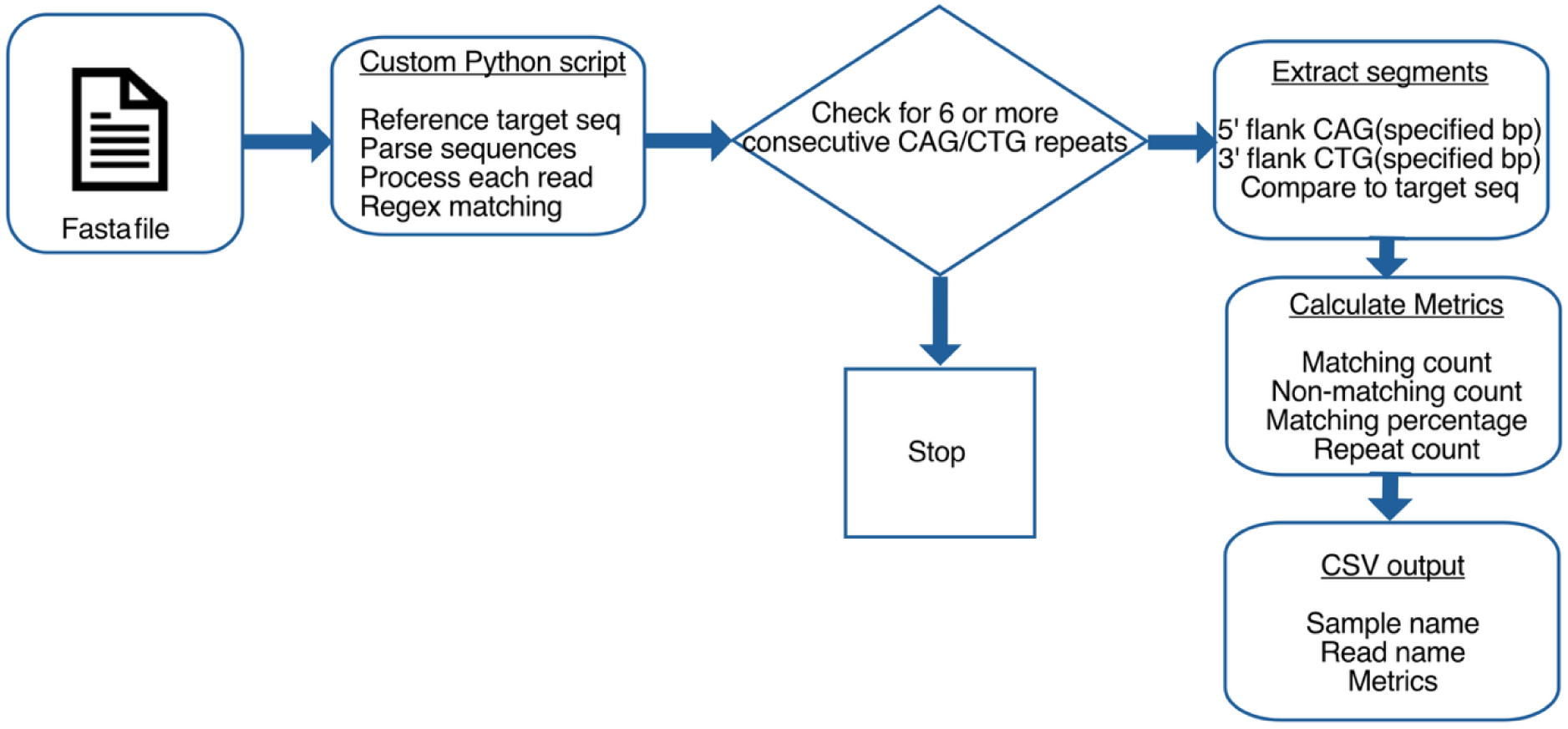
compare_sequence.py workflow to determine error frequencies in the sequence flanking the repeat tract.

